# An XA21-Associated Kinase (OsSERK2) regulates immunity mediated by the XA21 and XA3 immune receptors

**DOI:** 10.1101/000950

**Authors:** Xuewei Chen, Shimin Zuo, Benjamin Schwessinger, Mawsheng Chern, Patrick E. Canlas, Deling Ruan, Xiaogang Zhou, Jing Wang, Arsalan Daudi, Christopher J. Petzold, Joshua L. Heazlewood, Pamela C. Ronald

**Affiliations:** Department of Plant Pathology and the Genome Center, University of California, Davis, California 95616, USA; Rice Research Institute, Sichuan Agricultural University at Wenjiang, Chengdu, Sichuan 611130, China; Joint Bioenergy Institute, Emeryville, California 94710, USA; Key Laboratory of Crop Genetics and Physiology of Jiangsu Province, Key Laboratory of Plant Functional Genomics of the Ministry of Education, College of Agriculture, Yangzhou University, Yangzhou 225009, China

**Keywords:** immune receptor kinases, somatic embryogenesis receptor kinase (SERK), immunity, *Xanthomonas oryzae* pv. *oryzae*, Rice

## Abstract

The rice XA21 immune receptor kinase and the structurally related XA3 receptor, confer immunity to *Xanthomonas oryzae* pv. *oryzae* (*Xoo*), the causal agent of bacterial leaf blight. Here we report the isolation of OsSERK2 (rice somatic embryogenesis receptor kinase 2) and demonstrate that OsSERK2 positively regulates immunity mediated by XA21 and XA3 as well as the rice immune receptor FLS2 (OsFLS2). Rice plants silenced for *OsSerk2* display altered morphology and reduced sensitivity to the hormone brassinolide. OsSERK2 interacts with the intracellular domains of each immune receptor in the yeast-two-hybrid system in a kinase activity dependent manner. OsSERK2 undergoes bidirectional trans-phosphorylation with XA21 *in vitro* and forms a constitutive complex with XA21 *in vivo*. These results demonstrate an essential role for OsSERK2 in the function of three rice immune receptors and suggest that direct interaction with the rice immune receptors is critical for their function. Taken together, our findings suggest that the mechanism of OsSERK2-meditated regulation of rice XA21, XA3 and FLS2 differs from that of AtSERK3/BAK1-mediated regulation of Arabidopsis FLS2 and EFR.

## Introduction

The XA21 receptor kinase confers broad-spectrum resistance to *Xanthomonas oryzae* pv. *oryzae, (Xoo)* (Song et al., 1995). Animals and other plant species also carry membrane-anchored receptors with striking structural similarities to XA21 (Ronald and Beutler, 2010). Many of these receptors play key roles in recognition of conserved microbial signatures (also called pathogen-associated molecular patterns (PAMPs)) and host defense (Gomez-Gomez and Boller, 2000; Lemaitre et al., 1996; Medzhitov et al., 1997; Poltorak et al., 1998; Ronald and Beutler, 2010; Song et al., 1995; Zipfel et al., 2006). XA21 and structurally similar immune receptors activate defense signaling via membrane-associated complexes that include non-RD (arginine-aspartic acid) kinases to induce a core set of defense responses (Dardick et al., 2012; Ronald and Beutler, 2010). The non-RD kinases are either associated with the receptor via adaptor proteins (animals) or integral to the receptor (plants) (Ronald and Beutler, 2010; Schwessinger and Ronald, 2012). In rice the immune receptors XA21, XA3, PiD2 and FLS2, all belong to the non-RD subclass of kinases (Dardick et al., 2012; Schwessinger and Ronald, 2012).

In contrast to non-RD kinases, which are associated with the immune response, most RD kinases appear to regulate non-immune responses or serve as co-regulators of receptor kinase-mediated immunity (Chinchilla et al., 2006; Heese et al., 2007; Roux et al., 2011; Schwessinger and Ronald, 2012; Schwessinger et al., 2011) with the notable exception of the RD-kinase CERK1 in *Arabidopsis*, which directly binds chitin (Liu et al., 2012b).

In *Arabidopsis*, members of the somatic embryogenesis receptor kinase (SERK) regulate the function of multiple plasma membrane localized receptor kinases including hormone receptors and immune receptor kinases (Chinchilla et al., 2009; Li, 2010), and are members of the RD subclass of kinases (Schwessinger and Ronald, 2012). The best-studied member SERK3 is also referred to as BAK1 (brassinosteroid-insensitive 1 (BRI1) associated kinase 1) as it was initially identified as a key regulator of BRI1-mediated signaling (Li et al., 2002; Nam and Li, 2002). BRI1 is the main receptor of brassinosteroids (BR), an important class of plant hormones regulating growth and development (Clouse, 2011; Li and Chory, 1997). SERK3 and its closest paralog SERK4 are critical co-regulators of the immune response triggered by the ligand-activated *Arabidopsis* immune receptor kinase FLS2 (flagellin insensitive 2), EFR (EF-TU receptor), and PEPR1/2 (PEP receptors 1 & 2) (Chinchilla et al., 2006; Heese et al., 2007; Krol et al., 2010; Postel et al., 2009; Roux et al., 2011; Schwessinger et al., 2011). The pattern recognition receptors (PRRs) FLS2 and EFR recognize the bacterial proteins (or derived epitopes) flagellin (flg22) or EF-TU (elf18), respectively (Chinchilla et al., 2007; Zipfel et al., 2006). In contrast, PEPR1/2 are paralogous receptors for the endogenously produced small danger associated peptides, AtPeps (Yamaguchi et al., 2010; Yamaguchi et al., 2006). *Arabidopsis* FLS2 and EFR do not constitutively interact with SERK3 (or any other SERK family member) (Chinchilla et al., 2007; Heese et al., 2007; Roux et al., 2011; Schulze et al., 2010; Schwessinger et al., 2011). Only upon ligand binding, FLS2 and EFR undergo a nearly instantaneous complex formation with SERK3 and potentially with additional co-regulatory receptor kinases (Chinchilla et al., 2009; Roux et al., 2011; Schulze et al., 2010). The FLS2/EFR-SERK3 complex formation is independent of the kinase activity of either interaction partner or any other associated kinase (Schulze et al., 2010; Schwessinger et al., 2011). Indeed the co-crystal structure of the FLS2-SERK3 ectodomains and flg22 suggests that flg22 acts as a molecular glue by stabilizing the interaction between both receptors (Sun et al., 2013). This ligand-induced heteromer formation is the molecular switch-on for transmembrane signaling of these *Arabidopsis* receptor kinases (Albert et al., 2013). The tight association of the intracellular kinase domains is hypothesized to induce downstream signaling activation via specific structurally guided auto and transphosphorylation events. SERK3 and FLS2/EFR undergo unidirectional phosphorylation *in vitro*, that is, SERK3 is able to transphosphorylate FLS2 or EFR but not vice versa (Schwessinger et al., 2011).

In rice, XA21 confers robust resistance to *Xoo* (Song et al., 1995). XA21 biogenesis occurs in the endoplasmic reticulum (ER) (Park et al., 2010; Park et al., 2013). After processing and transit to the plasma membrane, XA21 binds to XB24 (XA21 binding Protein 24) (Chen et al., 2010b). XB24 physically associates with the XA21 juxtamembrane (JM) domain and catalyzes the autophosphorylation of serine and threonine residue(s) on XA21, keeping XA21 in an inactive state (Chen et al., 2010b). Upon pathogen recognition, XA21 kinase disassociates from XB24 and is activated (Chen et al., 2010b). This activation triggers a series of downstream events potentially including cleavage and nuclear localization of the XA21 kinase domain (Park and Ronald, 2012). XA21-mediated signaling is attenuated by the XB15 protein phosphatase 2C, which dephosphorylates XA21 (Park et al., 2008). Despite these advances, the early events governing XA21 activation have not yet been fully elucidated.

Based on the structural similarity of the XA3 immune receptor, which also confers immunity to *Xoo* (Sun et al., 2004; Xiang et al., 2006) and OsFLS2, which recognizes bacterial flagellin (Takai et al., 2008), with XA21, we hypothesized that XA3 and OsFLS2 transduce their responses through the same components that transduce the XA21-mediated response.

We have also identified an XA21 paralog lacking the transmembrane (TM) and kinase domains (called XA21D) (Wang et al., 1998). Based on the partial resistance phenotype conferred by XA21D and its predicted exclusively extracellular location, we hypothesized that XA21 and XA21D would partner with a co-regulatory receptor kinase (Wang et al., 1998). Based on recent findings (Chinchilla et al., 2007; Heese et al., 2007; Roux et al., 2011), we hypothesize that this hypothetical co-regulatory receptor kinase might be orthologous to *Arabidopsis* SERK proteins.

We therefore investigated the function of rice SERK family members in XA21-, XA3 and OsFLS2-mediated immunity. We isolated the RD receptor kinase OsSERK2 (Os04g38480) and demonstrated its requirement for both XA21- and XA3-mediated immunity as well as rice FLS2 signaling. We also show that OsSERK2 is involved in BR-regulated plant growth. The kinase domain of OsSERK2 directly interacts with XA3, XA21, OsFLS2 and OsBRI1 in yeast-two hybrid assays in an enzymatic activity dependent manner. Consistent with these results, OsSERK2 and XA21 form constitutive heteromeric complexes *in planta*. OsSERK2 and XA21 undergo bidirectional transphosphorylation *in vitro*, which is influenced by the domain architecture of both receptors. These results demonstrate an essential role for OsSERK2 in regulating development and receptor kinase-mediated immunity and suggest that direct interaction of OsSERK2 with the rice immune receptors is critical for function.

## Results

### Phylogenetic analysis of rice SERK family members

Previous studies in *Arabidopsis* demonstrated that SERK family members, and in particular SERK3 (also known as BAK1), are essential for both BR signaling mediated by BRI1 (Li et al., 2002; Nam and Li, 2002) and immunity mediated by FLS2, EFR and PEPR1/2 (Chinchilla et al., 2006; Heese et al., 2007; Krol et al., 2010; Roux et al., 2011; Schwessinger et al., 2011). In the case of rice XA21 and XA21D (an XA21 paralog lacking transmembrane and kinase domain), a co-regulatory receptor kinase has been hypothesized but its identification has remained elusive (Wang et al., 1998). Because XA21 is structurally similar to FLS2 and EFR and belongs to the same subfamily XII of LRR-RKs (Chen and Ronald, 2011), we hypothesized that one or more rice SERK family members serve as a co-regulatory receptor kinase for rice immune receptors.

To identify such a co-regulator, we carried out phylogenetic analysis on the two rice SERK proteins, OsSERK1 (Os08g07760) and OsSERK2 (Os04g38480) (Figure S1) (Singla et al., 2009), and five *Arabidopsis* SERK proteins (Li, 2010). The two rice SERKs are most closely related to *Arabidopsis* SERK1 and SERK2 (Figure 1A and Supplemental Figure 1). In contrast to *Arabidopsis* SERK3 (BAK1) and SERK4, which are the main SERK family members involved in *Arabidopsis* immune signaling (Li, 2010; Roux et al., 2011; Schwessinger et al., 2011), SERK1 and SERK2 are known for their role in developmental processes (Li, 2010). Recently it was shown that *Arabidopsis* SERK1 is also involved in immune signaling in transgenic plants expressing the tomato immune receptor Ve1 (Fradin et al., 2011). Nonspecific silencing of the two rice SERK proteins and several closely related proteins in rice compromises resistance against the fungal pathogen *Magnaporthe oryzae (M. oryzae*) (Park et al., 2011). Conversely, over-expression of OsSERK2 (referred to as OsSERK1 in the original publication (Hu et al., 2005)) enhances resistance against *M. oryzae* (Hu et al., 2005). These results suggest that one or both rice SERK proteins are involved in the rice immune response.

**Figure 1.**
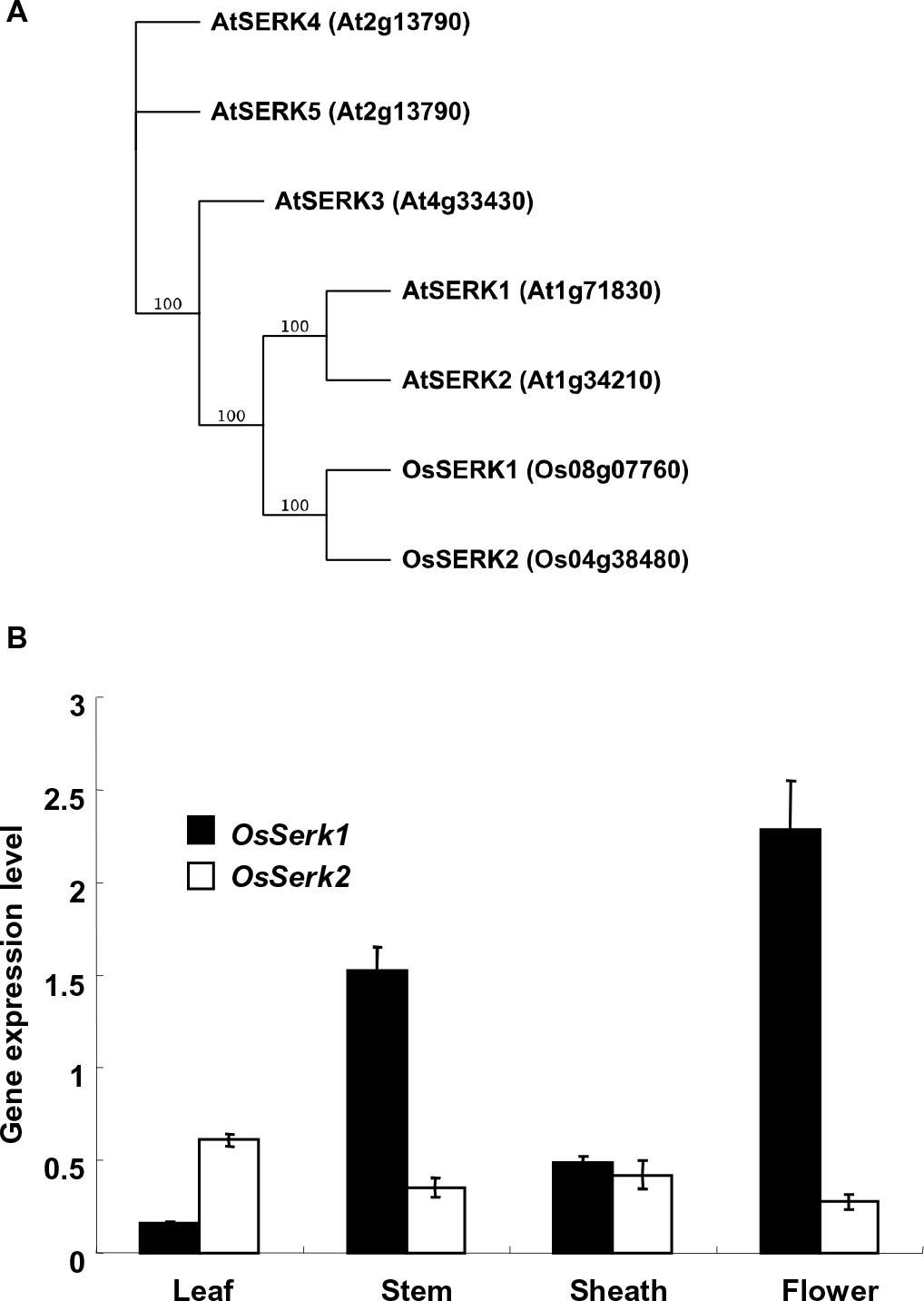
*OsSERK2* is the only rice *SERK*-family member highly expressed in leaf tissue. **(A)** Phylogenetic analysis of the two rice and five *Arabidopsis* SERK proteins. Rice SERK1 and SERK2 were grouped with their five *Arabidopsis* homologous SERK proteins. Full-length amino acid sequences of all SERK proteins were analyzed using Geneious Tree builder. The phylogenetic tree was generated using a bootstrap neighbor joining tree applying 1,000 replicates. Protein identifiers are given in brackets. **(B)** *OsSERK2* is the most highly expressed *SERK*-family member in mature rice leaves. Quantitative real-time PCR was performed on cDNA synthesized from RNA samples extracted from rice cultivar Nipponbare tissue as indicated. Gene expression levels of *OsSerk2* and *OsSerk1* were normalized to the expression of the *actin* reference gene. Data shown represent average expression level of one out three biological experiments with error bars indicating SD of three technical replicates. The experiment was repeated three times with similar results.

### *OsSerk2* is preferentially expressed in leaves whereas *OsSerk1* is expressed in flowers

Because *Xa21* confers resistance to *Xoo* in rice leaves (Song et al., 1995) we analyzed the expression patterns of *OsSerk1* and *OsSerk2* in leaves, stems, sheaths and flowers by performing quantitative RT-PCR. *OsSerk1* and *OsSerk2* were expressed in all tissues tested. *OsSerk2* is mainly expressed in leaves whereas *OsSerk1* is mainly expressed in flowers and stems (Figure 1B). The expression level of *OsSerk2* is much higher than that of *OsSerk1* in rice leaves (Figure 1B). These results suggest that *OsSerk2* rather than *OsSerk1* regulates *Xa21*-mediated immunity.

### Silencing of *OsSerk2* compromises Xa21-mediated immunity to *Xoo*

To test the function of OsSERK2 in rice XA21-mediated immunity, we carried out *OsSerk2* silencing experiments in the *Xa21* genetic background. For these experiments, we isolated a 383-bp *OsSerk2* cDNA fragment, which is unique to the *OsSerk2* gene, and introduced it into the pANDA vector, which carries a hygromycin selection marker, to generate the double-stranded RNA-based interference (dsRi) construct *pANDA OsSerk2Ri* (Miki and Shimamoto, 2004). We then introduced the *OsSerk2Ri* construct into *Xa21* and *ProA-tagged Xa21 (ProAXa21*) homozygous rice lines carrying the mannose selectable marker (Chen et al., 2010b). Three *Xa21-OsSerk2Ri* (abbreviated as *XOsSerk2Ri*) and one *ProAXa21-OsSerk2Ri* (abbreviated as *ProAXOsSerk2Ri*) double transgenic lines were assayed for silencing of *OsSerk2* using quantitative RT-PCR. Among these four double transgenic lines, two *XOsSerk2Ri* lines (*XOsSerk2Ri2* (*B-2*) and *XOsSerk2Ri3 (B-3)*) and the *ProAXOsSerk2Ri* line (*ProAXa21/X-B-1*) display specific reduction in the expression of *OsSerk2* (Supplemental Figure 2 and Supplemental Figure 3). The expression levels of *Xa21* and *OsSerk1* in these lines are similar to the control *Xa21* or *ProAXa21* lines (Supplemental Figure 2 and Supplemental Figure 3).

We then analyzed the response of the T_1_ plants from the double transgenic lines, *A814* derived from *B-2, A815* derived from *B-3* and *A804* derived from *ProAXa21/X-B-1*, to infection by the *Xoo* strain, PXO99AZ. Whereas the *Xa21* control line is highly resistant to *Xoo,* the double transgenic plants carrying *Xa21* and silenced for *OsSerk2* are susceptible, showing typical long water-soaked lesions (Figure 2). The susceptibility phenotype of the *OsSerk2* silenced lines co-segregates with the presence of the *OsSerk2Ri* transgene. Segregants from double transgenic lines carrying *Xa21* but lacking *OsSerk2Ri* are fully resistant (Supplemental Figure 4 and Supplemental Figure 5) demonstrating that silencing of *OsSerk2* compromises *Xa21*-mediated immunity in rice.

**Figure 2.**
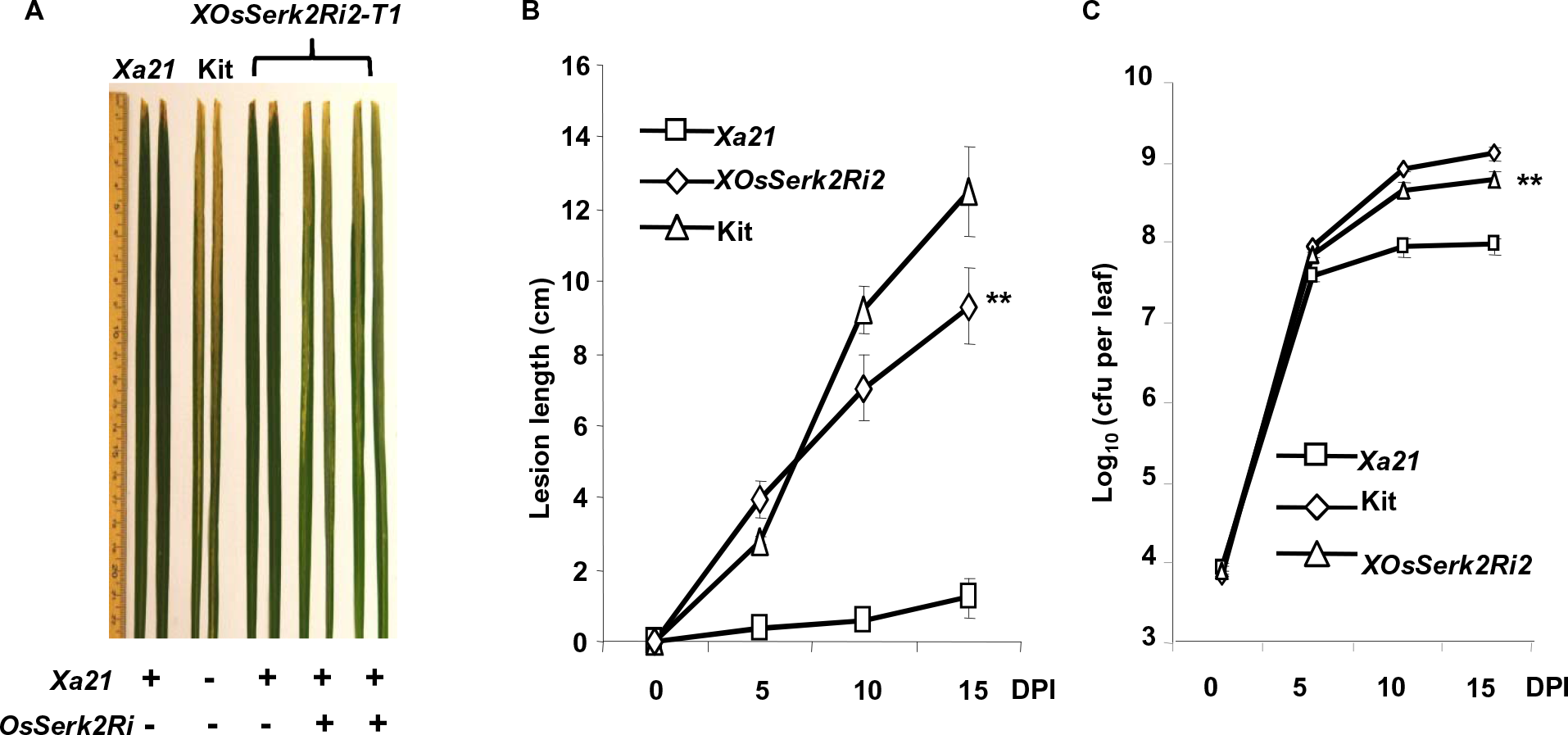
Silencing of *OsSerk2* compromises *Xa21*-mediated resistance to *Xoo* PXO99AZ. Six week-old plants of *Xa21-OsSerk2Ri-2* (*A814*), *Xa21* (resistant control) and Kitaake (Kit) (susceptible control) were inoculated with *Xoo* strain PXO99AZ. **(A)** *A814* plants in the presence of *OsSerk2Ri* develop long water-soaking lesions. Photograph depicts representative symptom development in leaves 14 days post inoculation. “+” and “-” indicate absence or presence of the *Xa21* and *OsSerk2Ri* transgene, respectively. **(B)** *XOsSerk2Ri2* plants (*A814-178*) homozygous for silenced *OsSerk2* develop long water soaked lesions. Lesion length was measured 0, 5, 10, and 15 days post inoculation. Graph shows average lesion length ± SD of at least 21 leaves from 7 independent plants. Statistical significance comparing *A814-178* with *Xa21* plants is indicated by asterisk (\*\**P* ≤ 0.05, ANOVA analysis, Tukey’s test). **(C)** *A814-178* is susceptible to *Xoo* PXO99AZ. Bacterial populations were counted 0, 5, 10, and 15 days post-inoculation. Each data point represents the average ± SD of six leaves from two independent plants. Statistical significance comparing *A814-178* with *Xa21* plants is indicated by asterisk (\*\**P* ≤ 0.05, ANOVA analysis, Tukey’s test). These experiments were repeated at least three times with similar results.

To further quantify the effect of *OsSerk2* silencing, we generated *Xa21* plants homozygous for the *OsSerk2Ri (A814-178*) transgene and performed more detailed infection studies. After 15 days of infection with *Xoo* PXO99AZ, the *A814-178* plants display long lesions, similar to the Kitaake control plants (Figure 2A). At 15 days post inoculation, the average lesion length (9.30 ± 1.04 cm) of *A814-178* plants is more than 7-fold greater than that of the *Xa21* plants (1.23 ± 0.55 cm). The observed lesion length difference between *A814-178* and *Xa21* plants is highly significant with a p-value less than 0.0003. The average disease lesion length of *A814-178* plants is closer to that of the susceptible parental control, Kitaake (12.5 ± 1.26 cm) (Figure 2B). Bacterial growth curve analysis revealed that the *Xoo* bacterial population in *A814-178* plants (6.51X10^8^±1.07X10^8^) is approximately 9-fold greater than in *Xa21* lines (7.47X10^7^±1.67X10^7^) and half of that observed for Kitaake (1.07X10^9^±2.01X10^8^) at 15 days post-inoculation (Figure 2C). These results are consistent with the leaf lesion phenotype described above. We also performed a similar experiment on an additional T_2_ homozygous double transgenic line (*A804-55*) developed from the *ProAXOsSerk2Ri* parent *A804* and obtained similar results (Supplemental Figure 6). These results demonstrate that silencing of *OsSerk2* compromises *Xa21*-mediated resistance. Using the same approach we silenced *OsSerk1* in the *Xa21-*Kitaake genetic background and analyzed the progeny for resistance. The silencing of *OsSerk1* did not affect *Xa21*-mediated immunity (Supplemental Table 1). These results indicate that *OsSerk2* but not *OsSerk1* is a key player in *Xa21*-mediated immunity.

### *OsSerk2* is essential for *Xa3*-mediated immunity

Like XA21, the rice XA3 resistance protein belongs to subfamily XII of the LRR receptor kinases, XA3 also functions as an immune receptor, conferring broad-spectrum resistance to most *Xoo* strains including PXO86 but not PXO99AZ (Sun et al., 2004; Xiang et al., 2006). Because of the structural and functional similarity of XA3 and XA21, we hypothesized that *OsSerk2* may also be required for *Xa3*-mediated innate immunity. To test this hypothesis we crossed *Xa3* plants (IRBB3) with the homozygous Kitaake *OsSerk2Ri-4 (Kit-B-4*) plants and obtained four F_1_ progeny called *Xa3OsSerk2Ri* F_1_ plants (Supplemental Figure 7). We inoculated these F_1_ plants with *Xoo* strain PXO86. As a control, we also inoculated the F_1_ progeny from a cross of *Xa3* and Kitaake plants. We found that F_1_ progeny carrying both *Xa3* and *OsSerk2Ri* displayed much longer lesions at 14 and 21 days post inoculation compared with F_1_ progeny of the control cross carrying *Xa3* but lacking *OsSerk2RiXa3* (ANOVA analysis: p-value less than 0.0001) (Figure 3A and 3B). To confirm the disease phenotype we monitored bacterial growth over time (Figure 3C). Fourteen days after inoculation, bacterial populations of *Xoo* strain PXO86 accumulated to nearly 100-fold higher levels in *Xa3OsSerk2Ri* plants when compared to F_1_ plants from the *Xa3*-Kitaake control cross (Figure 3C). These results show that OsSERK2 is also critical for resistance mediated by the XA3 immune receptor.

**Figure 3.**
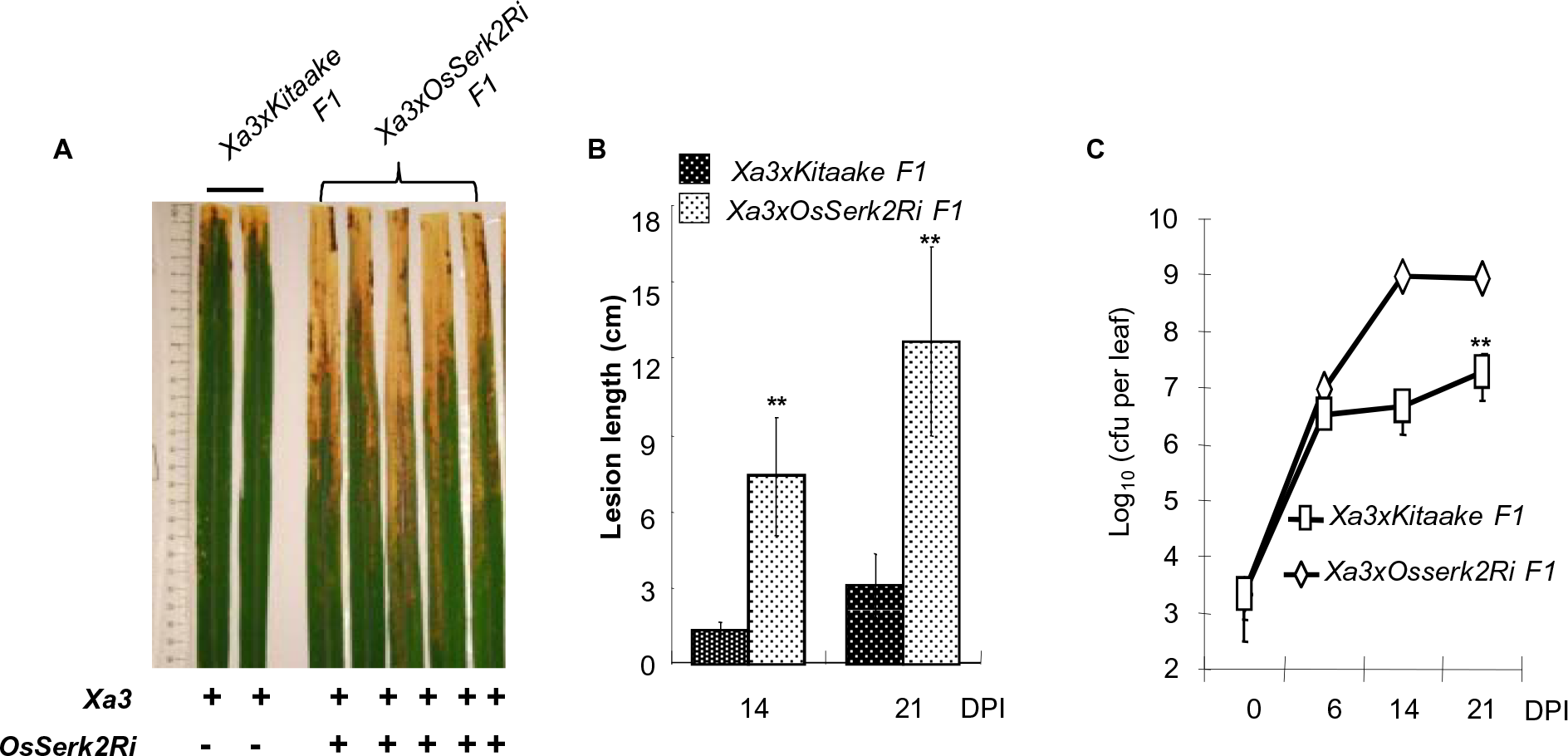
Silencing of *OsSerk2* compromises *Xa3*-mediated resistance to *Xoo* PXO86. Eight week-old plants of the F1 progeny of a cross between IRBB3 and Kitaake (*(Xa3/Kitaake)* F_1_, resistant control) and a cross between IRBB3 and Kitaake *OsSerk2RNAi(X-B-4-2)* homozygous for *OsSerk2RNAi (Xa3/X-B-4-2)* F_1_) were inoculated with *Xoo* strain PXO86. **(A)** (*Xa3/X-B-4-2*) F_1_ plants develop long water-soaking lesions. Photograph depicts representative symptom development in leaves 21 days post inoculation. “+” and “-” indicate absence or presence of the *Xa3* gene and *OsSerk2Ri* transgene, respectively. **(B)** (*Xa3/X-B-4-2*) F_1_ plants develop long water-soaked lesions. Lesion length was measured 14 and 21 days post inoculation. Graph shows average lesion length ± SD of at least 21 leaves from 7 independent plants. Statistical significance comparing (*Xa3/X-B-4-2*) F_1_plants with (*Xa3/Kitaake*) F_1_plants is indicated by asterisk (\*\**P* ≤ 0.05, ANOVA analysis, Tukey’s test). **(C)** (*Xa3/X-B-4-2*) F_1_ plants are susceptible to *Xoo* PXO86. Bacterial populations were counted 0, 6, 14, and 21 days post-inoculation. Each data point represents the average ± SD of six leaves from two independent plants from the same cross of *Xa3/X-B-4-2*. Statistical significance comparing (*Xa3/X-B-4-2*) F_1_ plants with (*Xa3/Kitaake*) F_1_ plants is indicated by asterisk (\*\**P* ≤ 0.05, ANOVA analysis, Tukey’s test). These experiments were repeated twice with similar results.

### OsSERK2 is involved in rice FLS2 mediated immune signaling

SERK3 and SERK4 associate with the PRR FLS2 *in vivo* and are important for FLS2-mediated signaling in *Arabidopsis* (Chinchilla et al., 2006; Heese et al., 2007; Roux et al., 2011; Schwessinger et al., 2011). Because flagellin is also able to trigger OsFLS2-mediated signaling in rice (Takai et al., 2008), we tested if OsSERK2 is involved in defense gene expression triggered by the application of flg22, a conserved peptide sequence derived from flagellin that is able to trigger FLS2-dependent defense signaling in many plant species including rice (Albert et al., 2010; Ding et al., 2012). We treated mature leaf strips of Kitaake or Kitaake plants silenced for *OsSerk2 (Kit-OsSerk2Ri-4*) (Supplemental Figure 7) with 1uM flg22 and measured the gene expression changes of two independent marker genes by quantitative RT-PCR. The expression of both *PR10b* and *Os4g10010* was dramatically reduced in plants silenced for *OsSerk2* (Figure 4A and 4B). Plants silenced for *OsSerk2* appeared to be fully sensitive to chitin application (Supplemental Figure 8) indicating that this reduction in defense gene expression was specific to flg22-triggered responses. These results suggest that chitin perception in rice is independent of OsSERK2 and is similar to SERK3-indpendent chitin perception in *Arabidopsis* (Chinchilla et al., 2009).

**Figure 4.**
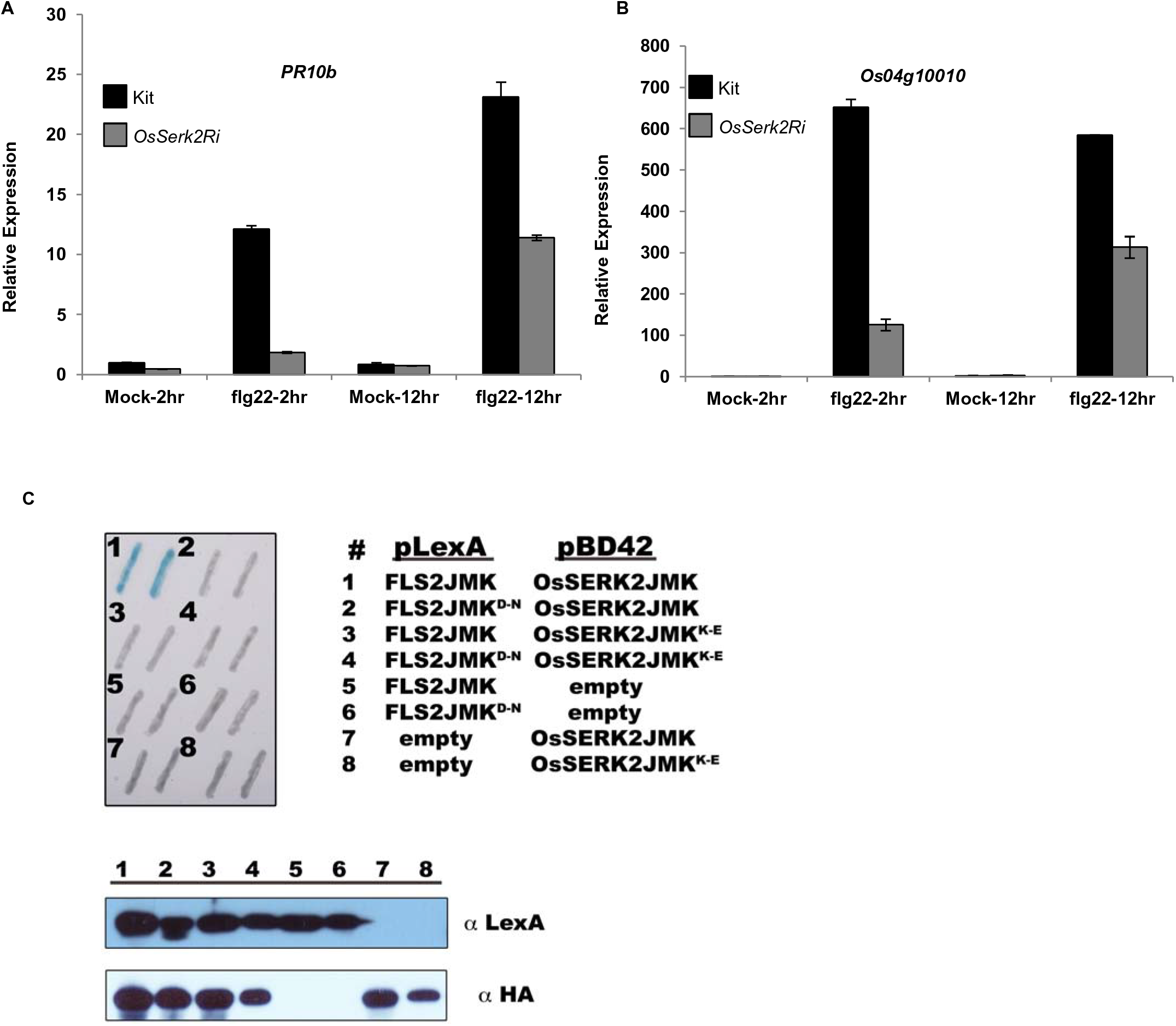
OsSERK2 regulates flg22-triggered defense gene expression in rice and directly interacts with the intracellular domain of FLS2 in the yeast-two hybrid system in a kinase catalytic activity dependent manner. Leaf strips of four-week old Kitaake control or *OsSerk2Ri* (*X-B-4-2*) plants were treated with 1 μM flg22 peptide for 2 or 12 hours. Expression levels of the two defense marker genes of *PR10b* **(A)** and *Os04g10010* **(B)** were measured by quantitative RT-PCR. Expression levels for each gene were normalized to *actin* reference gene expression. Data shown is normalized to the Kitaake mock treated (2 hour) sample. Bars depict average expression level ± SD of two technical replicates. This experiment was repeated four times with similar results. **(C)** OsFLS2 and OsSERK2 intracellular domains interact in a kinase dependent manner in the yeast-two hybrid system. Upper left panel: Two representative colonies for each co-transformation. The blue color indicates nuclear interaction between the two co-expressed proteins. Numbers indicate the specific co transformations. Upper right panel: Legend for the specific co-transformation events encoded by numbers. Lower panel: Western blot with anti-LexA or anti-HA antibodies to confirm expression of LexA and B42 fusion proteins, respectively, for each co-transformation event. The Matchmaker LexA two-hybrid system (Clontech) was used for the yeast two-hybrid assay.

Using the yeast two-hybrid system, we found that the intracellular domain of OsSERK2 interacts with the intracellular domain of OsFLS2 (#1 in Figure 4C) suggesting that OsSERK2 may directly regulate OsFLS2 function in rice. Mutations in residues required for full enzymatic activity in OsSERK2, OsFLS2 or both proteins compromised the interaction observed in the yeast-two hybrid systems (#2-4 in Figure 4C). This result suggests that full enzymatic activity of both OsFLS2 and OsSERK2 is required for complex formation.

### Rice plants silenced for *OsSerk2* display morphological features of BR-insensitive mutant plants and show reduced sensitivity to brassinolide

In *Arabidopsis* all four functional *SERK*-family members are involved in BRI1-mediated brassinosteroid signal transduction (Gou et al., 2012). We therefore hypothesized that OsSERK2 would regulate BR signaling in rice. Indeed, we found that *OsSerk2Ri* plants are semi-dwarf (Figure 5 A), similar to the *Osbri1* mutants plants (Nakamura et al., 2006). *XOsSerk2Ri2* plants are reduced in size compared with the *Xa21* control plants (Figure 5A). The leaf sheath, panicle and internodes of each tiller of *XOsSerk2Ri2* plants are shorter than those in *Xa21* control plants (Figure 5B, 5C and 5D). The lamina joint angle line is much reduced (2.8 ± 0.5^o^) compared to that of *Xa21* control plants (30 ± 3.2^o^) (P = 1.12X10^-29^, Student’s two-tailed T-test) (Figures 5E and 5F). The culm length of *XOsSerk2Ri2* is significantly shorter than that of *Xa21* control plants (Figure 5G). The relative lengths of internodes III and IV in the *XOsSerk2Ri2* plants are much reduced than those of *Xa21* control plants (Figure 5H). *OsSerk2Ri* plants exhibit shorter coleoptiles and show reduced sensitivity to brassinolide hormone (Supplemental Figure 9). These results demonstrate that OsSERK2 is also involved in rice BR hormone signaling.

**Figure 5.**
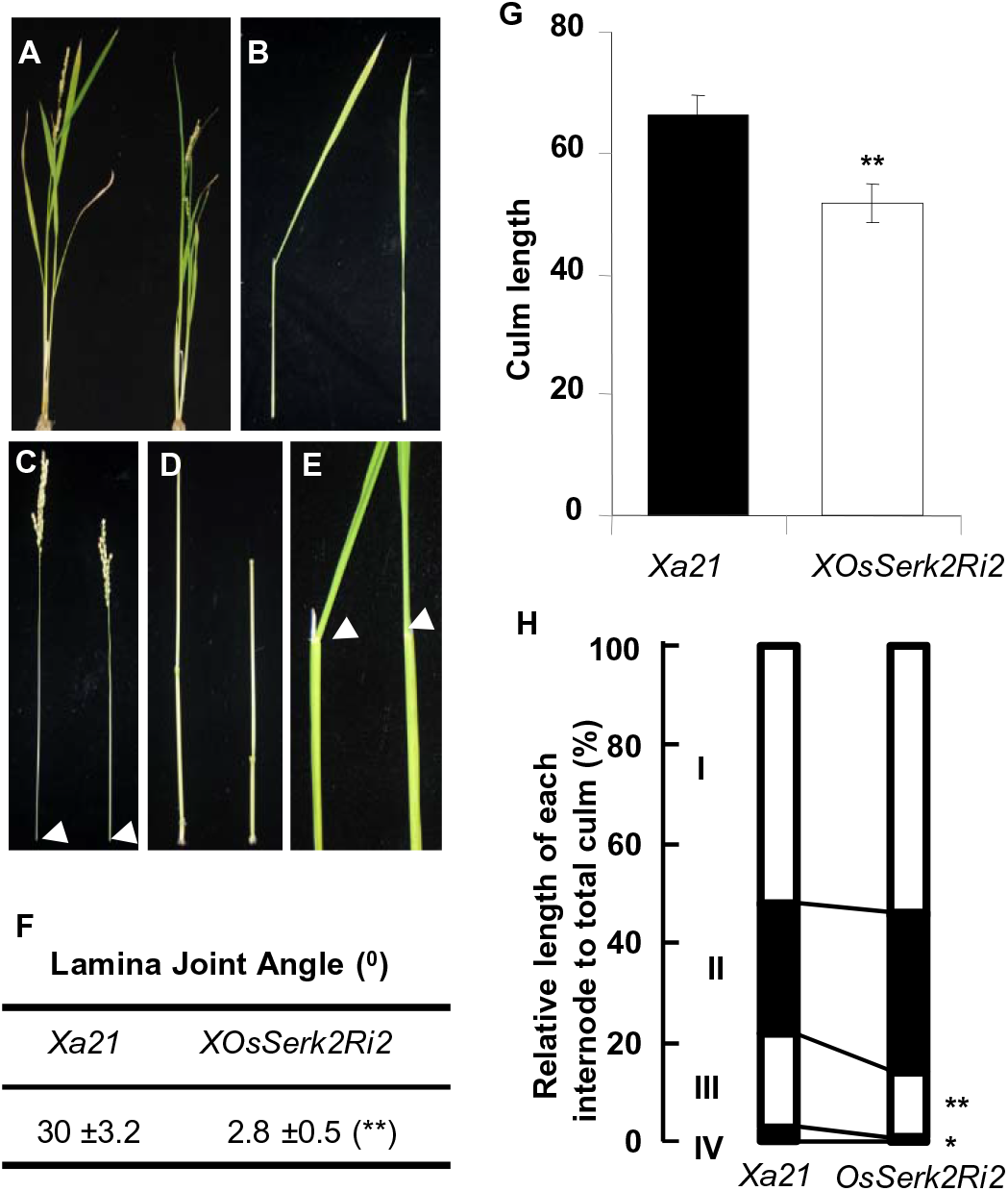
Plants silenced for*OsSerk2* display morphological features associated with compromised brassinosteroide signaling. **(A)** Gross morphology of *Xa21* control plants (left) and *A814-178* plants homozygous for silenced *OsSerk2* (right). **(B)** Leaf sheath morphology: the leaf sheath of *A814-178* (right) is shorter than in *Xa21* control plants (left). **(C)** Panicle structure: *A814-178* plants (right) have shorter panicle when compared to *Xa21* control (left) plants. The arrow heads indicate nodes. **(D)** Elongation pattern of internodes: the *Xa21* plants (left) show an N-type elongation pattern, whereas *A814-178* plants (right) show the typical dn-type pattern (Takeda, 1977). **(E)** Leaf morphology: leaves of *Xa21* control plants (left) are bent at the lamina joint indicated by the white arrowhead, whereas the leaves of *A814-178* plants (right) are erect. **(F)** Average degree of lamina joint angels of *Xa21* control and *A814-178* plants, respectively. (**G**) Measurement of the culm length from *Xa21* and *A814-178* plants, respectively. **(H)** Relative distance between internodes relative to total culm length in *Xa21* control and *A814-178* plants. In **(F)**, **(G)**,and **(H)**, the average ± SD of each parameter was determined from 12 plants of each genotype *Xa21* control and *A814-178*. (\**P* ≤ 0.1, \*\**P* ≤ 0.05, Student’s *t* test).

### OsSERK2 interacts with XA21, XA3 and OsBRI1 in a kinase-dependent manner in yeast

Next, we investigated if OsSERK2 directly regulates XA21-, XA3- and OsBRI1-mediated signaling. The rationale for this experiment was that it had previously been shown that SERK3 interacts with BRI1, EFR, FLS2 and PEPR1/2 (Li et al., 2002; Nam and Li, 2002; Postel et al., 2009; Roux et al., 2011). We performed yeast two-hybrid assays using XA21K668, a truncated version of XA21 containing the whole intra-cellular domain and part of the TM domain, which was previously shown to interact with several key XA21 binding proteins (Chen et al., 2010b; Park et al., 2008), as bait for interaction with OsSERK2. We found that OsSERK2JMK carrying part of the TM, as well as the JM and kinase (K) domain, interacts with XA21K668 (Figure 6A). Similarly, OsSERK2JMK is also able to interact with XA3JMK in the yeast-two-hybrid assay (Figure 6B). However, this interaction appears to be weaker than the interaction with XA21K668. OsSERK2JMK also interacts with OsBRI1JMK, the rice ortholog of *Arabidopsis* BRI1 (Figure 6C). In *Arabidopsis* the interactions between SERK3 and the ligand-binding receptor FLS2, EFR and BRI1 are independent of the catalytic activity of SERK3 when tested by *in planta* co-immunoprecipitation assays (Chinchilla et al., 2006; Schwessinger et al., 2011; Wang et al., 2008). To examine if this is also the case for OsSERK2, we tested the interaction between the catalytic inactive OsSERK2JMK^KE^ and XA21K668 or BRI1JMK in our yeast-two hybrid system. Both interactions were compromised by the catalytic inactivation of OsSERK2 (Figure 6A and 6C).

**Figure 6.**
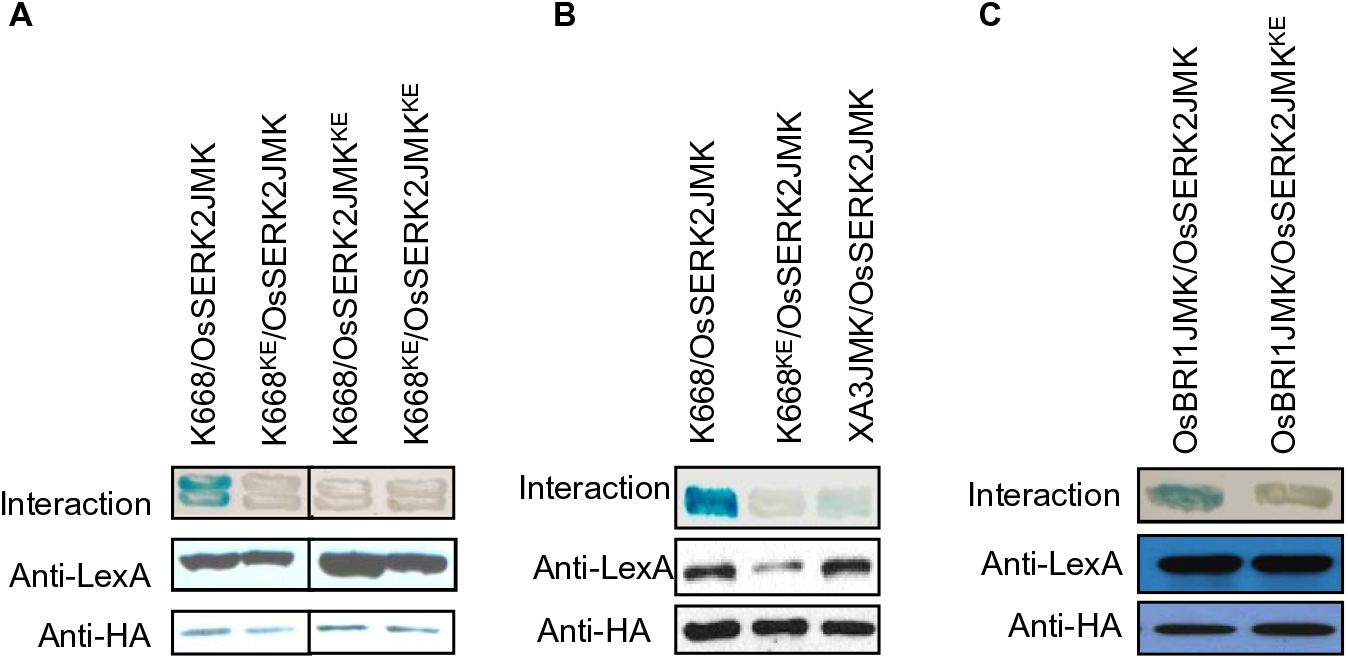
The OsSERK2intracellular domain interacts in a kinase activity dependent manner with the intracellular domain of the three predicted ligand-binding receptors XA21, XA3, and OsBRI1 in yeast-two hybrid system. The blue color indicates nuclear interaction between the two co-expressed proteins. (A) OsSERK2 and XA21 directly interact in a kinase catalytic activity dependent manner. Part of the transmembrane (TM) and the whole intracellular domain ofOsSERK2 (OsSERK2JMK) and its kinase catalytically inactive mutant OsSERK2JMK^K334E^ (OsSERK2JMK^KE^) were fused with the HA epitopeinthe vector pB42ADgc to obtain HA-OsSERK2JMK (abbreviated as OsSERK2JMK) and HA-OsSERK2JMK^KE^ (abbreviated as OsSERK2JMK^KE^). HA-OsSERK2JMK and HA-OsSERK2JMK^KE^ were co-transformed with LexA-XA21K668 (K668) or the catalytically inactive mutant LexA K668^K736E^ (abbreviated as K668^KE^), respectively. **(B)** OsSERK2 directly interacts with XA3. HA-OsSERK2JMK was co-transformed XA3, containing part of the TM and the whole intracellular domain used with LexA (LexA-XA3JMK (abbreviated as XA3JMK)). HA-OsSERK2JMK was also co-transformed with LexA-K668 and LexA-K668^KE^ for positive and negative interaction controls, respectively. **(C)** OsSERK2 and OsBRI1 directly interact in a kinase catalytic activity dependent manner. HA-OsSERK2JMK and HA-OsSERK2JMK^KE^ were co-transformed with OsBRI1 respectively, containing part of the transmembrane and the whole intracellular domain, fused with LexA (LexA OsBRI1JMK (abbreviated as OsBRI1JMK)). In (A), (B) and (C), the expression of LexA fused proteins, LexA-K668, LexA-K668^KE^, LexA-XA3JMK, and LexA-OsBRI1JMK was confirmed by Western blotting using an anti-LexA antibody. The expression of proteins, HA-OsSERK2JMK and HA-OsSERK2JMK^KE^, was confirmed by Western blotting using an anti-HA antibody. Yeast-two hybrid experiments were performed using Matchmaker LexA two-hybrid system (Clontech). This experiment was repeated three times with same results.

To assess if phosphorylation is critical for the interaction between XA21 and OsSERK2, we generated a suite of catalytically inactive protein variants including XA21JK, XA21JK^DN^, XA21K668^KE^, XA21K668^DN^, OsSERK2JMK^KE^, OsSERK2JK, OsSERK2JK^DN^ and OsSERK2TJK^DN^. The catalytically compromised protein variants where generated by either mutating the conserved lysine (K) required for ATP binding and catalytic activity or the aspartate (D) required for phospho-transfer (Nolen et al., 2004). We tested the interaction between these different protein variants in the yeast-two hybrid system (Figure 6A and Supplemental Figure 10). All mutant XA21 protein variants were compromised in the interaction with OsSERK2JMK and OsSERK2JK (Figure 6A and Supplemental Figure 10). Similarly no catalytically inactive protein variant of OsSERK2 was able to interact with XA21K688 (Supplemental Figure 9). Taken together, we conclude based on these results that the association of OsSERK2 with XA21, XA3 and OsBRI1 is dependent on the integrity of important catalytic residues and therefore most likely on the catalytic kinase activity of each protein in our yeast-two hybrid system.

### OsSERK2 forms a constitutive heteromeric complex with XA21 *in planta*

Next, we aimed to confirm the interaction between OsSERK2 and XA21 *in planta*. It was recently reported that the addition of fusion peptides to the carboxy terminus of *Arabidopsis* SERK3 interferes with its function in innate immune signaling (Ntoukakis et al., 2011). For this reason, instead of tagging OsSERK2, we raised an antibody (anti-OsSERK2) against a unique peptide consisting of 10 amino acids (602-611) at the C-terminus of OsSERK2. Initially, we tested the specificity of anti-OsSERK2 using the *E.coli*-produced GST-OsSERK2JMK protein. Because the protein encoded by *OsSerk1* is the closest paralog to OsSERK2, we included the *E.coli*-produced GST-OsSERK1JMK protein in our experiment as a control. The anti-OsSERK2 antibody specifically recognizes OsSERK2 but not OsSERK1 (Figure 7A). The presence of two bands corresponding to GST-OsSERK2JMK suggests that GST-OsSERK2JMK is strongly phosphorylated during heterologous protein production in *E. coli* (Wang et al., 2005). Indeed, when we treated GST-OsSERK2JMK with the highly active lambda-phosphatase the upper bands corresponding to (hyper-)phosphorylated GST-OsSERK2JMK disappeared and GST-OsSERK2JMK was detected as a discrete band of a single molecular size (Supplemental Figure 11). We also tested the specificity of the anti-OsSERK2 antibody on total protein extracted from *Xa21* rice plants and *Xa21* rice plants silenced for *OsSerk2 (XOsSerk2Ri2*). The anti-OsSERK2 antibody specifically recognized a protein band of the approximate size of 70 kDa very close to the molecular mass of OsSERK2. This band was only present in the total protein extract of *Xa21* plants but not in the total protein extracts of plants silenced for *OsSerk2 (XOsSerk2Ri2*) (Figure 7B and Supplemental Figure 12). This observation suggests that the anti-OsSERK2 antibody specifically recognizes the OsSERK2 protein in total rice protein extracts.

**Figure 7.**
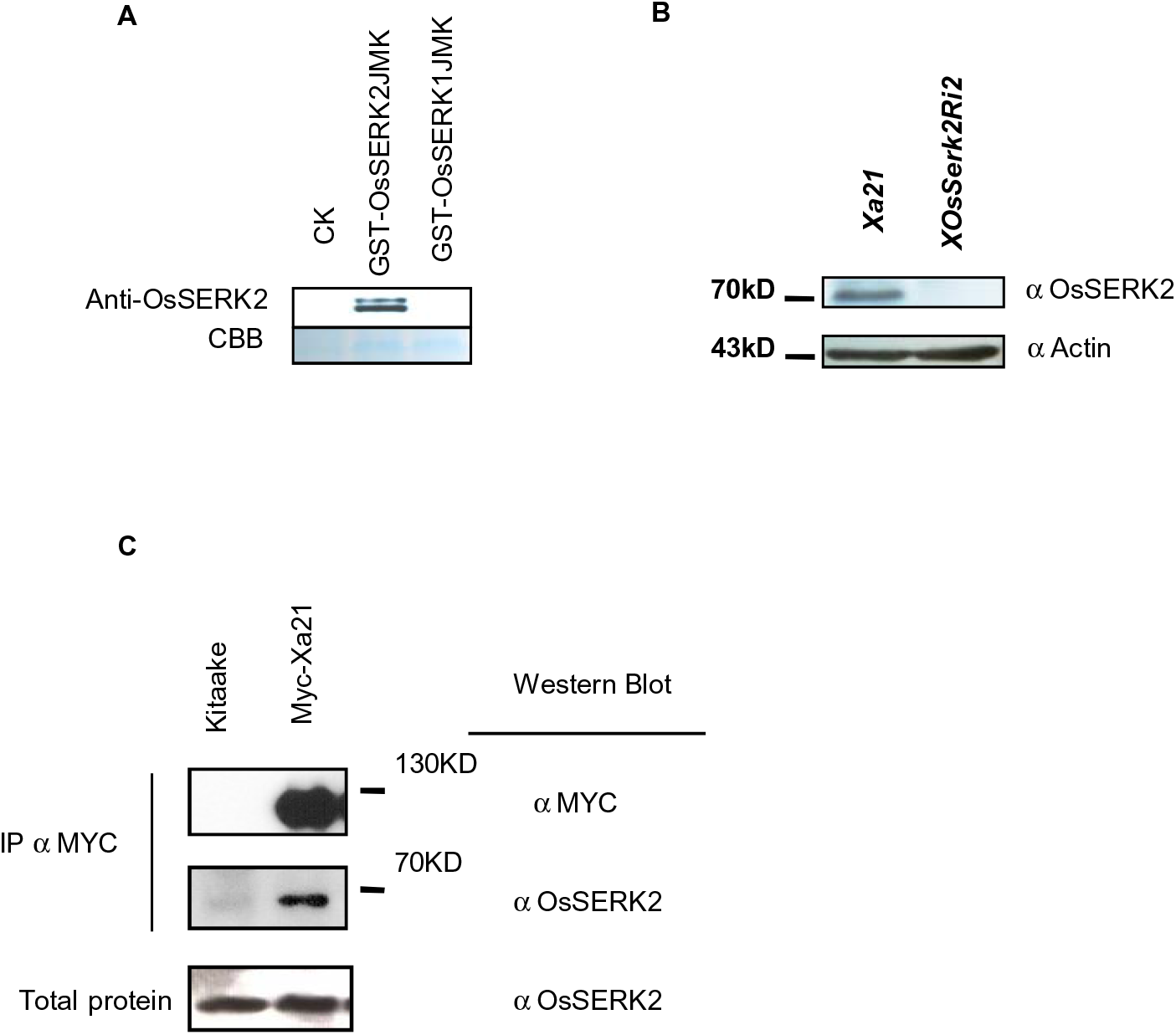
OsSERK2 and XA21 form constitutive complexes *in planta*. **(A)** The newly developed anti-OsSERK2 antibody raised against a specific 10 amino acid epitope at the carboxy terminus specifically recognizes OsSERK2 but not OsSERK1. Top panel shows an anti-OsSERK2 western blot on in *E.coli* expressed and purified GST-OsSERK1JMK, GST-OsSERK2JMK and GST-control proteins. The lower panel shows the Coomassie Brilliant Blue (CBB) staining of the corresponding region to assess equal quantities of protein was loaded. **(B)** The anti-OsSERK2 antibody recognizes a specific protein of approximately 70 kDa in size in total rice protein extracts from *Xa21* plants but not from *OsSerk2*-silenced *Xa21* plants (*XOsSerkRi2*) (upper panel). 75ug of total protein for each genotype were separated by SDS-PAGE gel electrophoresis and subjected to immuno-blot analysis with anti-OsSERK2 antibody (upper panel) or anti-Actin antibody (lower panel) as loading control. Full membranes for each immune-blot are shown in Supplemental Figure 12. **(C)** OsSERK2 and XA21 form constitutive ligand-independent complexes *in vivo.* Immuno-complexes were precipitated from leaf material of *Myc-Xa21* expressing rice plants using agarose-conjugated anti-Myc antibody. Kitaake rice leaves were used for the negative control. Components of the immuno-precipitated complexes were separated by SDS-PAGE gel followed by immuno-detection with anti-Myc (for Myc-XA21) and anti-OsSERK2 (for OsSERK2), separately. Myc-XA21 gives a band at about 130 kDa. OsSERK2 (～70KD) was co-immunoprecipitated with XA21 in the absence of any treatment. The lower panel shows equal amounts of OsSERK2 in both total protein fractions before immunoprecipitation. This experiment was repeated four times with similar results.

To further investigate the association between XA21 and OsSERK2 *in vivo*, we carried out co-immunoprecipitation experiments on protein extracts from mature leaves of 4-5 week old greenhouse grown plants at a stage when XA21 signaling is fully functional and *Xoo* infection assays are usually performed (Century et al., 1999). We previously generated transgenic plants carrying fully functional Myc-*Xa21* under the control of its native promoter (Park et al., 2008). In co-immunoprecipitation experiments with anti-Myc conjugated agarose beads we detected a ～130 kDa polypeptide using the anti-Myc antibody only in transgenic plants (Figure 7C, top panel). This is consistent with our previous reports (Chen et al., 2010b; Park et al., 2008; Park and Ronald, 2012).

We next used the anti-OsSERK2 antibody to test if OsSERK2 is present in the immunoprecipitated complexes. We successfully detected a band corresponding to the predicted size of OsSERK2 of ～70 kDa in plants expressing Myc-XA21 but not in the wild-type Kitaake control (Figure 7C, middle panel). OsSERK2 was readily detected in the mock control sample indicating that XA21 and OsSERK2 can be found in constitutive complexes *in planta* (Figure 7C, middle panel).

### OsSERK2 and XA21 undergo bidirectional transphosphorylation events *in vitro* depending on their domain architecture

As both OsSERK2 and XA21 are receptor kinases, we tested whether transphosphorylation occurs between these two kinases. For this purpose, we expressed and purified a suite of GST-or His-NUS-tagged truncated protein variants of OsSERK2 and XA21, respectively (Figure 8A). We first investigated the phosphorylation capacities of XA21K668 and OsSERK2JMK both containing part of the TM, full JM and kinase domains (Figure 8A). Either kinase incubated on its own in the presence of [^32^P]-γ-ATP is able to undergo autophosphorylation (Figure 8B, 8C and Supplemental Figure 13). This phosphorylation is abolished by mutations in the ATP-binding site demonstrating that the observed effect was not due to a co-purified kinase (Figure 8B, 8C and Supplemental Figure 13). Next we incubated each kinase with a catalytically inactive counterpart. Using protein variants in which both kinases contain part of the TM domain, we found that OsSERK2JMK was able to transphosphorylate XA21K668 but not the reverse (Figure 8B). The inability of XA21K668 to transphosphorylate OsSERK2 is independent of the domain structure of OsSERK2 or of the residue mutated to compromise catalytic activity (Figure 8B). XA21K668 is also unable to transphoshorylate OsSERK2JK variants that lack the TM domain and consist exclusively of the entire intracellular domain (Supplemental Figure 13).

**Figure 8.**
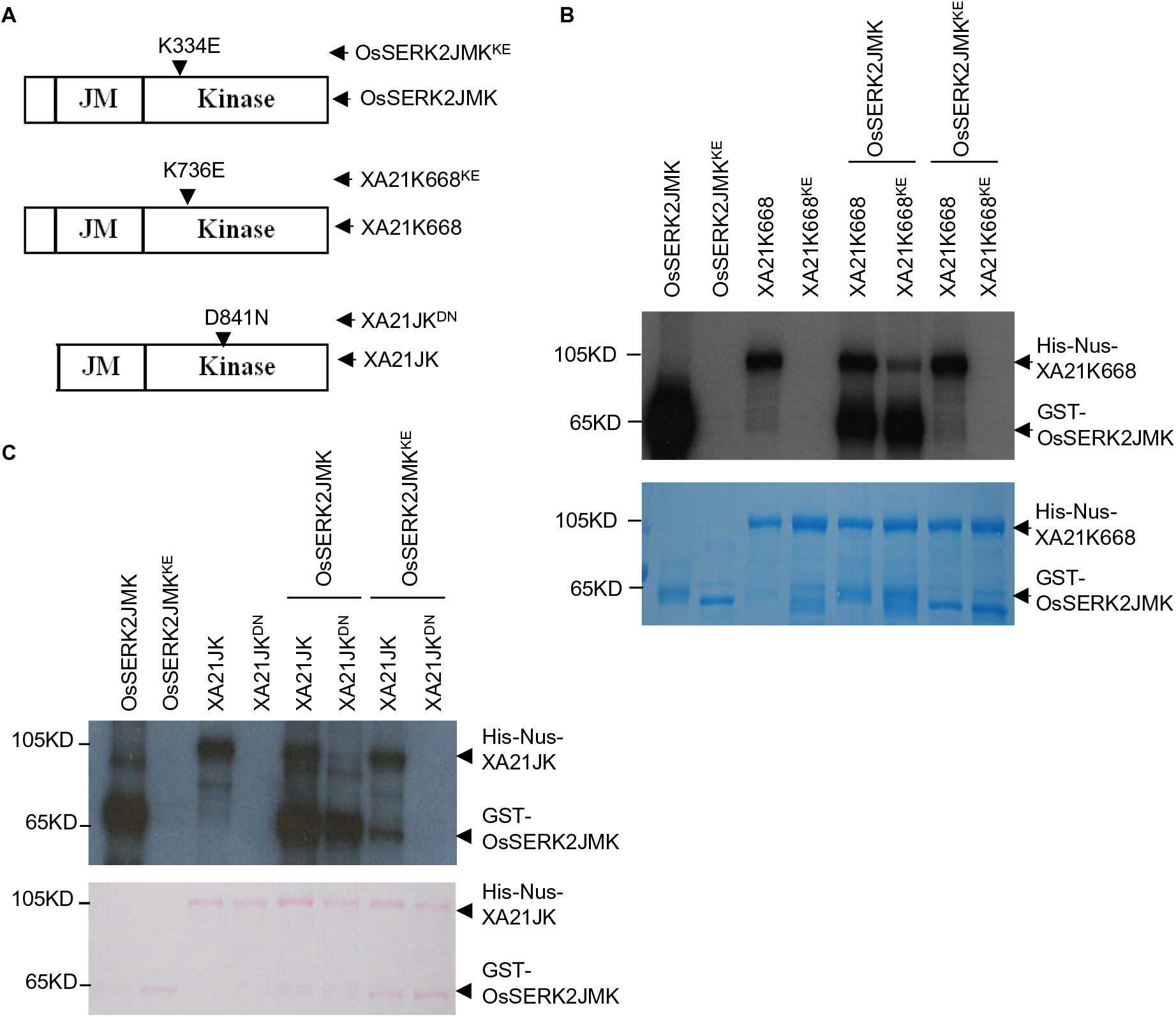
OsSERK2 and XA21 undergo bidirectional transphosphorylation depending on their domain architecture *in vitro*. **(A)** Depiction of protein domain architecture used for the trans-phosphorylation assays. OsSERK2JMK, XA21K668 and their respective kinase inactive variants, OsSERK2JMK^K334E^ (OsSERK2JMK^KE^), XA21K668^K736E^ (XA21K668^KE^) proteins contain partial sequences of their TM domain and full juxtamembrane (JM) and kinase domains. XA21JK, and its kinase inactive variant XA21JK^D841N^ (XA21JK^DN^) contain the full JM and kinase domain but lack the partial TM domain. **(B)** OsSERK2JMK is able to trans-phosphorylate XA21K668 but not *vice versa*. The assay was performed by incubating GST-OsSERK2JMK (abbreviated as OsSERK2JMK) and GST-OsSERK2JMK^K334E^ (abbreviated as OsSERK2JMK^KE^) in the presence or absence of His-Nus-XA21K668 (abbreviated as XA21K668), and His-Nus-XA21K668 and His-Nus-XA21K668^K736E^ (abbreviated as XA21K668^KE^) in the presence or absence of GST-OsSERK2JMK using radioactive labeled [^32^P]-γ-ATP. Proteins were separated by SDS/PAGE and analyzed by autoradiography in the top panel and the protein loading control by CBB staining is shown in the lower panel, respectively. **(C)** XA21JK is able to transphosphorylate OsSERK2JMK but not vice versa. The assay was performed by incubating GST-OsSERK2JMK (abbreviated as OsSERK2JMK) and GST-OsSERK2JMK^K334E^ (abbreviated as OsSERK2JMK^KE^) in presence or absence of His-Nus-XA21JK (abbreviated as XA21JK), and His-Nus-XA21JK and His-Nus-XA21JK^D841N^ (abbreviated as XA21JK^DN^) in the presence or absence of GST-OsSERK2JMK using radioactive labeled [^32^P]-γ-ATP. Proteins were separated by SDS/PAGE and analyzed by autoradiography in the top panel and the protein loading control is shown by Ponceau S in lower panel, respectively. This experiment was repeated twice with similar results.

Next we tested the catalytic capacity of XA21JK, a XA21 protein variant exclusively consisting of the entire intracellular domain (Figure 8A). In the absence of any TM domain XA21JK is able to transphosphorylate the catalytic inactive version of OsSERK2JMK (Figure 8C). In this set-up OsSERK2JMK is unable to phosphorylate XA21JK^D841N^ (Figure 8C).

These observations suggest that XA21 and OsSERK2 undergo bidirectional phosphorylation *in vitro*. In addition, the capacity of XA21 kinase to function as phospho-group-acceptor and donor is influenced by the presence of the TM domain. A similar observation has been recently made for the human epidermal growth factor receptor (EGFR) (Wang et al., 2011).

### OsSERK2 is an active kinase that undergoes autophosphorylation at multiple serine and threonine residues *in vitro*

We previously demonstrated that XA21 exhibits relatively low autophosphorylation activity *in vitro* and is mostly autophosphorylated in the juxtamembrane region, which is important for its function (Chen et al., 2010a; Chen et al., 2010b). In contrast, OsSERK2 exhibits a much stronger autophosphorylation activity (Figure 8 and Supplemental Figure 13) and potentially undergoes multiple autophosphorylation events similar to its *Arabidopsis* ortholog (Karlova et al., 2006; Oh et al., 2010; Wang et al., 2008). To identify authophoshorylation sites of OsSERK2, we performed mass spectrometry on OsSERK2JK after incubation with cold ATP. We identified twelve unique phosphorylation events on serine and threonine residues (Table 1, Supplemental Figure 14 and Supplemental Data 1). The phosphorylation sites are evenly distributed over the entire intracellular domain of OsSERK2. Comparison with previously published phosphorylation sites of *Arabidopsis* SERK1 to SERK3 (Karlova et al., 2006; Oh et al., 2010; Wang et al., 2008) revealed that the *in vitro* phosphorylation pattern of OsSERK2 is most closely related to AtSERK1 rather than AtSERK3 (Table 1), further validating the phylogenetic analysis (Figure 1). Overall, the phosphorylation sites within the activation segments are conserved between all SERK proteins (residues T459, T463, T464 and T468 in OsSERK2). In contrast residues predicted to be involved in protein-protein interactions and downstream signaling (all other residues of OsSERK2) appear to be specific to each individual SERK protein (Table 1).

**Table 1:**
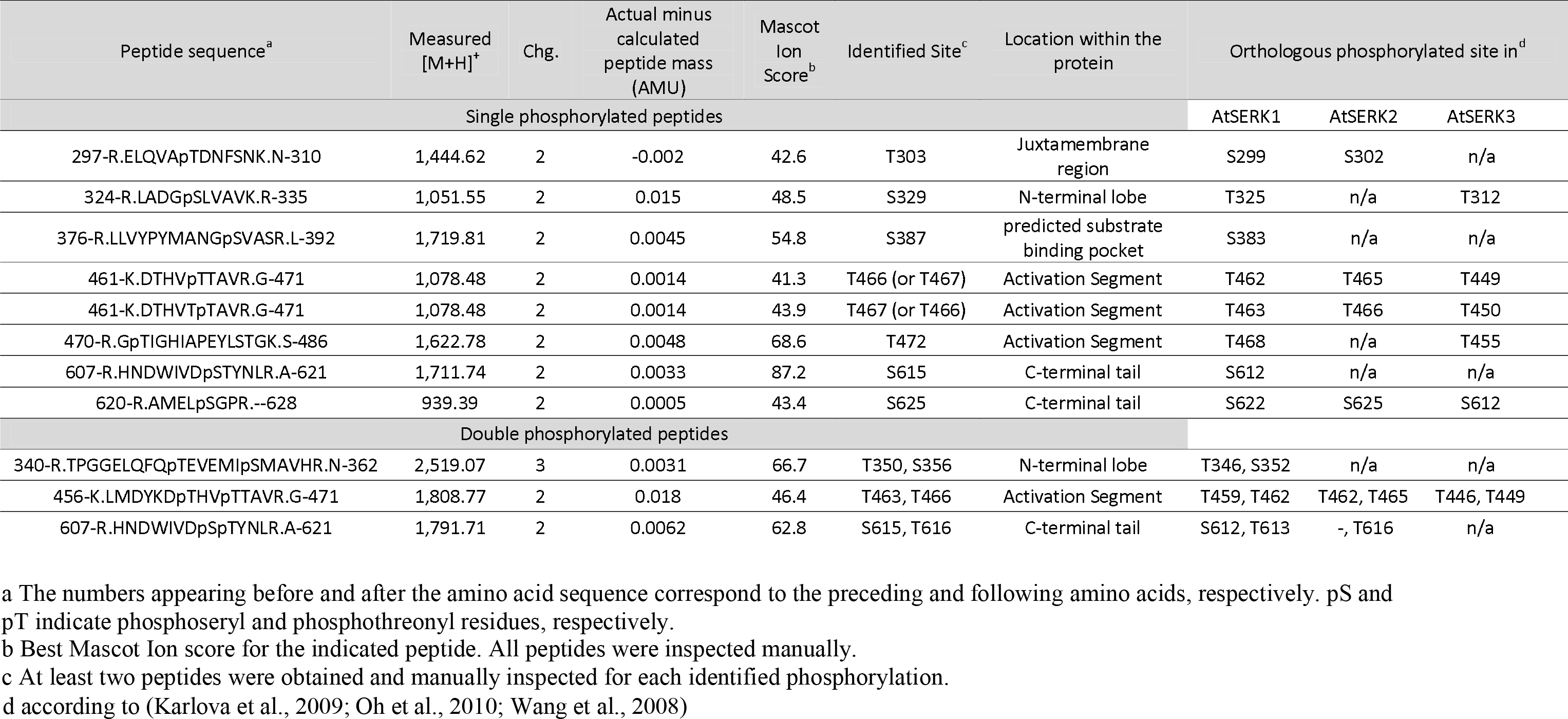
Identification of *in vitro* OsSERK2 phosphorylation Sites by Q-ToF LC/MS/MS and their conservation in Arabidopsis SERK1, SERK2 and SERK3. For each identified phosphorylation site the highest scoring peptide, its specific parameters and its conservation in three Arabidopsis SERK proteins is given. The specific MS2 spectra can be found in Supplementary Figure 12 and in Supplementary Data 1.

## Discussion

It was previously reported that OsSERK2 is involved in BR signaling and in resistance against the fungal pathogen *M. oryzae* (Hu et al., 2005; Park et al., 2011). Silencing of *OsSerk2* in combination with *OsSerk1* and other related genes leads to stunted growth, reduced sensitivity to exogenous BR application and compromised resistance to *M. oryzae* (Park et al., 2011). In these studies, it was consistently shown that over-expression of *OsSerk2* enhances resistance to *M. oryzae* (Hu et al., 2005). A detailed molecular mechanism of how the altered expression of *OsSerk2* leads to these phenotypes has not been previously provided. Here we show that OsSERK2 is required for immune signaling pathways controlled by three immune receptor kinases: XA21, XA3 and OsFLS2 (Figure 2, 3, 4, Supplemental Figure 4, 5, and 6), but is not required for CeBIP-mediated chitin signaling (Supplemental Figure 8). In addition, we conclusively demonstrate that *OsSerk2* is required for BR signaling in rice (Figure 5 and Supplemental Figure 9). Because OsSERK2 interacts with the intracellular domain of these immune receptors in the yeast-two hybrid system, OsSERK2 most likely exerts its regulatory function by directly interacting with and phosphorylating these receptors (Figure 4, 5 and Supplemental Figure 10). In yeast, the interaction between XA21 and OsSERK2 requires the catalytic activity of both kinases (Figure 5A and Supplemental Figure 10).

These observations suggest that the catalytic activity of each interaction partner is required for formation of stable constitutive heteromeric complexes between XA21 and OsSERK2. In addition, if the catalytic activity of either kinase is compromised these proteins might still be able to transiently interact as shown by the transphosphorylation assays between active and catalytic impaired kinases *in vitro* (Figure 8 and Supplemental Figure 10). This transient interaction between XA21 and OsSERK2 might explain how a catalytic impaired variant of XA21 is able to confer a partial resistance phenotype (Andaya and Ronald, 2003). Several of the newly identified autophosphorylation sites of OsSERK2 might be important for a stable interaction with XA21 and downstream signaling (Table 1). In future studies it will be interesting to analyze the contribution of individual phosphorylation sites of OsSERK2 on its role in BR and immune receptor kinase signaling in rice.

### Multiple functional roles of rice OsSERK2

Our domain structure and phylogenetic analysis indicates that the rice genome encodes only two SERK proteins: OsSERK1 and OsSERK2 (Figure 1A, Supplemental Figure 1, Supplemental Figure 14A and 14B). Although previous reports hypothesized the presence of several additional SERK-like proteins (Li et al., 2009; Singla et al., 2009), our analysis shows that these additional candidates lack at least one of the characteristic structural features of SERK-proteins: five extracellular LRR domains, a proline rich region, a transmembrane domain or an intracellular kinase domain (Supplemental Figure 15A and 15B) (Hecht et al., 2001). In addition only OsSERK1 and OsSERK2 cluster with the five *Arabidopsis* SERKs whereas the next ten closest rice homologs that contain five extracellular LRR domains do not (Supplemental Figure 15A and 15B).

In *Arabidopsis*, the five SERK proteins are involved in diverse signaling pathways and are often functionally redundant (Li, 2010). SERK1, SERK2, SERK3 and SERK4 interact with BRI1 and function as positive regulators of BL signaling (Albrecht et al., 2008; Gou et al., 2012; Karlova et al., 2006; Li et al., 2002; Nam and Li, 2002). SERK1 and SERK2 play redundant roles in male sporogenesis (Albrecht et al., 2005). SERK1 has recently been shown to be involved in organ separation in flowers (Lewis et al., 2010). Both SERK3 and SERK4 regulate cell death and senescence (He et al., 2007; Kemmerling et al., 2007). Importantly, SERK3 and SERK4 are also both required for FLS2-, EFR-, and PEPR1/2-mediated innate immune responses (Roux et al., 2011; Schwessinger et al., 2011).

These observations suggest that the SERK proteins in *Arabidopsis* may have undergone functional diversification/specification. The fact that the over-expression of the two rice SERK-like proteins Os02g18320 and Os06g12120 is able to partially rescue the BR insensitive phenotype of the *bri1-5* mutation in *Arabidopsis* suggests that rice SERK-like proteins (Supplemental Figure 15A and 15B) can also fulfill functions previously only attributed to SERK proteins (Li et al., 2009).

In *Arabidopsis* SERK3 and SERK4 are required for innate immunity to biotrophic, hemibiotrophic and necotrophic pathogens (Kemmerling et al., 2007; Roux et al., 2011; Schwessinger et al., 2011). In contrast, only OsSERK2, but not OsSERK1, significantly contributes to rice immunity to the biotrophic bacterial pathogen *Xoo* (Figure 2–4 and Supplemental Figure 4-6) and the hemibiotrophic fungal pathogen *M. oryzae* (Hu et al., 2005; Park et al., 2011). OsSERK2 may mediate its immunity against *M. oryzae* through a yet-to-be characterized immune receptor. Suppression of *OsSerk2* expression in transgenic calli by RNA interference results in a significant reduction in the rate of shoot regeneration indicating that OsSERK2 is a positive regulator of somatic embryogenesis in rice (Hu et al., 2005). In contrast, over-expression of *OsSerk2* increases the rate of shoot regeneration (Hu et al., 2005). OsSERK2 is also involved in BR hormone signaling in rice (Figure 5, 6C and Supplemental Figure 9). *OsSerk2*-silenced rice plants show a similar morphology as the *Osbri1* mutant (Nakamura et al., 2006) (Figure 5), are less sensitive to exogenous BR application (Supplemental Figure 8) and OsSERK2 directly interacts with OsBRI1 (Figure 6C). The fact that transgenic over-expression of *OsSerk2* in the *Arabidopsis bri1-5* mutant partially rescues its BR insensitive phenotype (Li et al., 2009) also supports that OsSERK2 functions in BR hormone signaling. It is clear that OsSERK2 functions in signaling pathways regulating multiple developmental programs and rice innate immune responses. How OsSERK2 regulates these multiple signaling pathways and if these pathways are cross-co-regulated remains to be determined. Recent studies investigating the crosstalk between BR mediated growth and innate immune signaling in *Arabidopsis* reached conflicting conclusions on the requirement of SERK3 (Albrecht et al., 2012; Belkhadir et al., 2012).

OsSERK2 is phylogenetically most closely related to *Arabidopsis* SERK1 and SERK2 (Figure 1) and its *in vitro* autophosphorylation pattern is closest to that of SERK1 (Table 1) but not SERK3 and SERK4. It would be informative to test if OsSERK2 is able to complement immune related phenotypes of SERK3 mutants in *Arabidopsis* and if any of the differential autophosphorylation sites are involved in this process. These experiments might determine whether the phylogenetic diversification of the *SERK* gene family in *Arabidopsis* is driven by functional specification of certain family members and their specific phosphorylation pattern.

### Mechanistic differences between rice and *Arabidopsis* in the perception of conserved microbial signatures

Here we report the functional conservation of one rice SERK protein in innate immune signaling mediated by three pattern recognition receptors. SERK proteins are involved in the immune response towards a plethora of distinct pathogens in multiple plant species including *Nicotiana benthamiana*, *Solanum lycopersicum*, *Lactuca sativa* L., *Oryza sativa* and *Arabidopsis thaliana* (Chaparro-Garcia et al., 2011; Fradin et al., 2009; Hu et al., 2005; Kemmerling et al., 2007; Mantelin et al., 2011; Park et al., 2011; Roux et al., 2011; Santos and Aragao, 2009; Schwessinger et al., 2011). The molecular mechanism of how SERK proteins exert their function in these immune pathways is well-studied in the only case of SERK3/BAK1 in *Arabidopsis*. SERK3/BAK1 was shown to interact with multiple pattern recognition receptors such as FLS2 and EFR in a ligand-dependent and kinase independent manner *in planta* using whole 2-week-old seedlings grown in sterile media, in sterile *Arabidopsis* cell-cultures or when transiently overexpressed in *N. benthamiana* leaves (Roux et al., 2011; Schulze et al., 2010; Schwessinger et al., 2011). This heteromeric complex formation was shown to be quasi-instantaneous in the case of SERK3/BAK1 and FLS2 in *Arabidopsis* cell-cultures (Schulze et al., 2010) suggesting that these proteins constitutively co-localize in plasma-membrane subdomains ready to signal. Recent crystallographic studies show that flg22 serves as a molecular glue between FLS2 and BAK1 and stabilizes the complex between both ectodomains (Sun et al., 2013). This ligand-dependent rapid heteromeric complex formation is thought to be a key molecular switch for activating plant immune receptor-mediated signaling in *Arabidopsis* (Albert et al., 2013).

In rice, the interaction between OsSERK2 and XA21 occurs in the absence of any ligand treatment in fully mature leaves of 4-5 week old greenhouse grown plants (Figure 7B) indicating a comparatively strong constitutive heteromeric complex formation. Another mechanistic difference between rice and *Arabidopsis* is that it appears that in rice the complex formation between OsSERK2 and distinct PRRs, XA21 and FLS2, requires the catalytic activity of each interacting kinase (at least in the yeast-two hybrid assay) (Figure 4C, 6 and Supplemental Figure 10). To our knowledge, direct interaction between the kinase domains of *Arabidopsis* SERK proteins and FLS2 or EFR in the yeast-two hybrid system have not been reported to date. The observed ability of the kinase domains of rice SERK proteins and their respective rice PRR counterparts to form constitutive heteromeric complexes in yeast in the absence of the ligand (or any ectodomain) could be a further indication that the interaction is more strongly mediated by their kinase domains in rice when compared to *Arabidopsis*.

The mechanistic differences of microbial signature perception in rice and *Arabidopsis* are not restricted to the interaction between OsSERK2, XA21 and other PRRs. Several recent reports demonstrate the differential involvement of homologous proteins in chitin and peptidoglycan perception (PGN) when comparing rice and *Arabidopsis*. For example, both plant species utilize LysM-containing proteins in chitin and PGN perception. However, the mechanism with which these proteins directly bind to the corresponding conserved microbial signatures to trigger signal transduction is clearly distinct. In rice the GPI-anchored LysM-containing proteins CEBIP, LYP4 and LYP6 directly bind to chitin and the latter two also bind bacterial PGNs (Kaku et al., 2006; Liu et al., 2012a). The rice LysM receptor-like kinase CERK1 forms direct heteromeric complexes with CeBIP and is required for chitin signaling but is most likely not involved in direct chitin binding (Shimizu et al., 2010). Therefore rice CERK1 appears to be a downstream receptor-like kinase relying the extracellular chitin (and potentially PGN) perception into an intracellular defense response.

In contrast, in *Arabidopsis* CERK1 is the major chitin binding protein and is required for chitin perception (Miya et al., 2007; Petutschnig et al., 2010; Wan et al., 2008). Even though *Arabidopsis* CEBIP is able to biochemically bind chitin it is not involved in chitin perception (Shinya et al., 2012). This clearly suggests that in *Arabidopsis* CERK1 is the sole functional chitin receptor. In addition CERK1 is also required for PGN perception in *Arabidopsis* but does not directly bind to PGN (Willmann et al., 2011). Instead *Arabidopsis* LYM1 and LYM3, the two closest homologs of rice LYP4 and LYP6, specifically bind to and are required for PGN signaling but not chitin (Willmann et al., 2011).

These examples of chitin and PGN perception and the data presented in this study demonstrate that homologous proteins are involved in the perception of conserved microbial signatures in rice and *Arabidopsis*. Yet the molecular mechanisms and the specific involvement of each protein can be distinct.

### Model for XA21 signal transduction from the plasma membrane to the nucleus

Our *in vitro* phosphorylation assays show that OsSERK2 directly transphosphorylates XA21 only if it contains parts of its TM domain. In this structural configuration, XA21 is unable to phosphorylate OsSERK2 (Figure 8 and Supplemental Figure 13). In contrast, the XA21 truncated protein that contains the complete intracellular domain, but lacks the TM domain (XA21JK), is capable of transphosphorylating OsSERK2 (Figure 8 and Supplemental Figure 13). These results have several implications. First, they demonstrate that XA21 has the capacity to transphosphorylate OsSERK2 under the appropriate conditions, but not under conditions where XA21 contains part of the TM domain. Second, they suggest a possible mechanism by which XA21 is activated and transduces the signal from the plasma membrane to the nucleus: the XA21 kinase is kept inactive by structural features mediated by its TM domain. In this scenario, ligand binding to the XA21 extracellular domain would induce conformational changes in the XA21-OsSERK2 complex and subsequently trigger transphosphorylation of XA21 by OsSERK2. Downstream signaling might be in part mediated by nuclear localization of the activated, cleaved XA21 kinase domain (Park and Ronald, 2012). In addition OsSERK2 might phosphorylate downstream signaling components such as BIK1-like kinases at the plasma membrane (Lu et al., 2010; Zhang et al., 2010).

## Materials and Methods

### Plant growth, *Xoo* inoculation and disease resistance determination

Transgenic *Xa21*, *cMycXa21*, *ProAXa21* plants were generated in the Kitaake genetic background (Chen et al., 2010b; Park et al., 2008). Rice IRBB3 carrying the LRR receptor kinase XA3 (Sun et al., 2004; Xiang et al., 2006) was used for *Xa3*-related experiments. The rice Nipponbare genetic background was used for analysis of the transcriptional expression of *OsSerk1* and *OsSerk2*. All transgenic plants in Kitaake were grown in the greenhouse until 6 weeks of age and transferred to the growth chamber before *Xoo* (PXO99AZ) inoculation. IRBB3 plants, which carry the endogenous *Xa3* and confer resistance to *Xoo* strain PXO86 (Sun et al., 2004; Xiang et al., 2006), were grown in the greenhouse until two months of age and transferred to the growth chamber before *Xoo* (PXO86) inoculation. In the green house, the light intensity in photosynthetic photon flux across the spectrum from 400 to 700 nm was approximately 250 μmol m^−2^ s^−1^ in spring. The growth chamber was set on a 14-hour daytime period, a 28/26°C temperature cycle and at 90% humidity. The chamber was equipped with metal halide and incandescent lights. The light intensity in the growth chamber was approximately 100 μmol m^−2^ s^−1^. Bacterial suspensions (OD_600_ of 0.5) of *Xoo* were used to inoculate rice by the scissors-dip method (Song et al., 1995). The disease lesion length and bacterial population accumulated in rice leaf were evaluated as reported previously (Chern et al., 2005). Statistical analysis was performed using the appropriate statistical analyses.

### Generation of rice transgenic plants and F1 progeny

RNAi constructs *OsSerk2Ri* and *OsSerk1Ri* were introduced into *Xa21*, *ProAXa21* or Kitaake plants through *Agrobacterium*-mediated transformation according to the method described previously (Chern et al., 2005). Because the *Xa21* and *ProAxa21* transgenic plants are mannose resistant, transgenes *OsSerk2Ri* and *OsSerk1Ri* were selected with hygromycin in our studies. The plants of transgenic line *X-B-4-2* homozygous for *OsSerk2Ri* in the Kitaake genetic background (abbreviated as *OsSerk2Ri*) were used for crossing with IRBB3 to obtain *Xa3OsSerk2Ri* plants. The cross was performed using IRBB3 as the pollen donor. PCR-based genotyping on *Xa21*, *OsSerk2Ri* and *OsSerk1Ri* was performed as described previously (Chen et al., 2010b).

### RNA extraction and quantitative RT-PCR analyses

Total RNA was isolated from rice plant tissues using TRIzol (Invitrogen), following the manufacturer’s instructions. Total RNA was treated with DNase I (NEB) and used for first strand cDNA synthesis using the Invitrogen reverse transcription kit (Invitrogen) following the provided manual. Quantitative real time PCR (qRT-PCR) was performed on a Bio-Rad CFX96 Real-Time System coupled to a C1000 Thermal Cycler (Bio-Rad). For qRT-PCR reactions, the Bio-Rad SsoFast EvaGreen Supermix was used. qRT-PCR primer pairs used were as follows: *OsSerk2-Q1*/*Q2* (5’-TAGTCTGCGCCAAAGTCTGA -3’/5’-GCACCTGACAGTTGTGCATT -3’) for the *OsSerk2* gene, *OsSerk1*-*Q1*/-*Q2*(5’-TGCATTGCATAGCTTGAGGA -3’/5’- GCAGCATTCCCAAGATCAAC -3’) for the *OsSerk1* gene, *Xa21*-*Q1*/-*Q2*(5’-TGACACGAAGCTCATTTTGG-3’/5’-TTGATGGCATTCAGTTCGTC-3’) for the *Xa21* gene, *Os04g10010-Q1/-Q2* (5’-AAATGATTTGGGACCAGTCG*-*3’/5’- GATGGAATGTCCTCGCAAAC*-*3’) for *Os04g10010* gene, *PR10b-Q1/-Q2 (*5’- GTCGCGGTGTCGGTGGAGAG*-*3’, 5’-ACGGCGTCGATGAATCCGGC*-*3’) for *PR10b* gene, Actin*-Q1*/*-Q2* (5’-TCGGCTCTGAATGTACCTCCTA-3’/ 5’-CACTTGAGTAAAGACTGTCACTTG-3’) for the reference gene *actin*. qRT-PCR reactions were run for 40 cycles with annealing at 62^0^C for 5 sec and denaturation at 95^0^C for 5 sec. The expression levels of *OsSerk2*, *OsSerk1, Os04g10010, PR10b* and *Xa21* were normalized to the *actin* gene expression level.

### Constructions

All constructs were made according to supplemental experimental procedures (Text S1).

### Purification of recombinant proteins and kinase assay

Purification of GST- or His-Nus- fusion proteins and *in vitro* kinase and transphosphorylation assays were performed as described previously (Liu et al., 2002; Schwessinger et al., 2011).

### Defense gene expression analysis

Fully developed leaves of 6 week old rice plants were cut into 2 cm long strips and incubated for at least 12 hours in ddH20 to reduce residual wound signal. Leaf strips were treated with 1μM flg22 peptide (Felix et al., 1999), purchased from Pacific Immunology, or 50 ug/mL chitin, purchased from Sigma, for the indicated time. Leaf tissue was snap-frozen in liquid nitrogen and processed as described above.

### Yeast Two-Hybrid Assays

The Matchmaker LexA two-hybrid system (Clontech) was used for yeast two hybrid assays. Yeast pEGY48/p8op-lacZ (Clontech) was co-transformed with the BD and AD vectors by using the Frozen-EZ yeast transformation II kit (Zymo Research) and spread on an appropriate medium following the procedures described previously (Chen et al., 2010b).

### Immunoblotting

Total protein extraction from yeast, *E. coli*, and rice plants and immuno-blotting (Western blotting) were performed as previously described (Chen et al., 2010b). The anti-OsSERK2 antibody against the synthetic peptide AELAPRHNDW-Cys of OsSERK2 (amino acids 602-611) was provided as a service by Pacific Immunology. Detailed information about their methods can be obtained at Pacific Immunology (http://www.pacificimmunology.com/). Anti-OsSERK2 for detection of OsSERK2, anti-LexA (Clontech) for detection of LexA-fused protein produced from BD vectors, anti-HA (Covance) for detection of HA-fused protein produced from AD vectors, and anti-Myc (Santa Cruze Biotech) for detection of XA21 with Myc tag were used as primary antibodies.

### Co-Immunoprecipitation of rice proteins

Detached rice leaves from six weeks-old *cMyc-Xa21* or Kit plants were cut into 4 cm long pieces and snap frozen. Myc-XA21 complex was immunoprecipitated using the agarose conjugated anti-Myc antibody (Santa Cruz) following the method described previously (Roux et al., 2011; Schwessinger et al., 2011) with slight adaptation. The immunoprecipitates were then probed with anti-Myc and anti-OsSERK2, respectively, after being separated by SDS-PAGE.

### Brassinolide (BL) treatment

Seeds from *Xa21*, *A814-178*, and *A814-186* were sterilized with 30% bleach for 20 mins. After rinsing with distilled ddH2O four times, they were germinated in a growth chamber at 30°C on MS agar in the presence or absence of 0 µM, 0.001 µM, 0.01 µM, and 0.1 µM of 24-epiBL (Sigma). Seedlings were examined 5 days after germination.

### Phylogenetic and molecular evolutionary analyses

Phylogenetic and molecular evolutionary analyses were conducted using MUSCLE in the Geneious (Biomatters) environment. 1000 bootstraps were adopted to infer the statistical support for the tree.

### Tandem Mass Spectrometry (LC-MS/MS)

Samples were analyzed on an Agilent 6550 iFunnel Q-TOF mass spectrometer (Agilent Technologies) coupled to an Agilent 1290 LC system (Agilent). Peptide samples were loaded onto a Ascentis Peptides ES-C18 column (2.1 mm x 100 mm, 2.7 μm particle size; Sigma-Aldrich, St. Louis, MO) via an Infinity Autosampler (Agilent Technologies) with buffer A (2% Acetonitrile, 0.1% Formic Acid) flowing at 0.400 mL/min. Peptides were eluted into the mass spectrometer via a gradient with initial starting conditions of 5% buffer B (98% acetonitrile, 0.1% formic acid) increasing to 35% B over 5.5 minutes. Subsequently, B was increased to 90% over 30 seconds and held for 3 minutes at a flow rate of 0.6 ml/min followed by a ramp back down to 5% B over one minute where it was held for 2.5 minutes to re-equilibrate the column. Peptides were introduced to the mass spectrometer from the LC by using a Jet Stream source (Agilent Technologies) operating in positive-ion mode (5000 V). The data were acquired with the Agilent MassHunter Workstation Software, LC/MS Data Acquisition B.05.00 (Build 5.0.5042.2) operating in Auto MS/MS mode whereby the five most intense ions (charge states 2 to 5) within a 300 to 1400 m/z mass range above a threshold of 1000 counts were selected for MS/MS analysis. MS/MS spectra were collected with the quadrupole set to “Medium” resolution and collision energy dependent on the m/z to optimize fragmentation (3.6 x (m/z) / 100–4.8). MS/MS spectra were scanned from 100 to 1700 m/z and were acquired until 45000 total counts were collected or for a maximum accumulation time of 333 ms. Former parent ions were excluded for 0.1 minutes following selection for MS/MS acquisition.

### Analysis of tandem mass spectrometry data

Mass spectral data were initially examined in the Agilent MassHunter Workstation Software, Qualitative Analysis B.05.00 (Build 5.0.519.13 Service Pack 1). All MSMS data were exported in MGF format from the Qualitative Analysis software using; Absolute height >= 20 counts, Relative height >= 0.100% of largest peak, Maximum number of peaks (limited by height) to the largest 300, Peak spacing tolerance 0.0025 m/z plus 7.0 ppm, Isotope model: Peptides, Limit of assigned charge states to a maximum of 5. Resultant. mgf files were used to interrogate the Mascot search engine version 2.3.02 (Matrix Science) with a peptide tolerance of ± 20 ppm and MS/MS tolerance of ± 0.1 Da; variable modifications were Oxidation (M), Phospho (ST), Phospho (Y); up to one missed cleavage for trypsin; require bold red; and the instrument type was set to ESI-QUAD-TOF. Searches were performed against the current protein set (all.pep, release 7.0) from the MSU Rice Genome Annotation Project (Kawahara et al., 2013) including standard contaminants (keratin, trypsin, GFP, BSA etc.) resulting in a database with 66,506 sequences and 29,649,083 residues. An initial Iions score or expect cut-off of 20 was applied to filter low-scoring phosphopeptide matches. All phosphopeptide matches were manually inspected and annotated to confirm the modification. A minimum of two independent spectra were inspected for each phosphorylation site.

## Acknowledgments

This work was supported by NIH GM59962 to PCR. Dr. X C was also supported by National Natural Science Foundation of China (NSFC 31171622), Sichuan “Hundred Talents Plan” fund and Sichuan Agricultural University “High Talents” start-up fund in China. B.S. was supported by an EMBO long-term fellowship. B.S. is supported by a Human Frontiers Science Program fellowship. We are grateful to Dr. Wenming Wang (Sichuan Agricultural University, Chengdu, Sichuan, China) for his helpful discussion on this manuscript.

## Author contributions

Conceived and designed the experiments: X.C., B.S., S.Z. and P.C.R. Performed the experiments: X.C., B.S., S.Z., P. E. C., D. R., X. Z, A. D., C. P., J. H. and J.W. Analyzed the data: B.S., X.C, S.Z., M.C., C. P., J. H. and P.C.R. Wrote the paper: B.S., X.C., M.C. and P.C.R.

The authors declare no conflict of interests.

---

**Table S1. Disease lesion length determination on T_1_ plants of the *OsSerk1Ri* rice**

“Expression of *OsSerk1*” represents the expression level of *OsSerk1* in *Xa21-OsSerk1Ri (XOsSerk1Ri*) and *ProAXa21-OsSerk1Ri (ProAXOsSerk1Ri*)T_0_ transgenic lines. T_1_ plants were genotyped using a primer pair targeting the *hygromycin* resistance gene (primer sequence needed). “Hyg(+)” indicates transgenic plants carrying *OsSerk1Ri*.

